# Phase-specific pooling of sparse assembly activity by respiration-related brain oscillations

**DOI:** 10.1101/2021.06.09.447658

**Authors:** Shani Folschweiller, Jonas-Frederic Sauer

## Abstract

Nasal breathing affects cognitive functions, but it has remained largely unclear how respiration-driven inputs shape information processing in neuronal circuits. Current theories emphasize the role of neuronal assemblies, coalitions of transiently active pyramidal cells, as the core unit of cortical network computations. Here, we show that respiration-related oscillations (RROs) directly pace the activation of neuronal assemblies in the medial prefrontal cortex (mPFC) of mice. Neuronal assemblies are more efficiently entrained than single neurons and activate preferentially during the descending phase of RROs. At the same time, overlap between individual assemblies is minimized during descending RRO due to the efficient recruitment of GABAergic neurons by assemblies. Our results thus suggest the RROs support cortical operations by defining time windows of enhanced yet segregated assembly activity.

## Introduction

Nasal airflow activates olfactory sensory neurons in the olfactory epithelium (Grosmaitre et al., 2007), thereby producing oscillating depolarizations that are broadcast to the brain via the olfactory bulbs (Fontanini and Bower, 2006). Besides the well-studied role of respiration-related oscillations (RROs) in the processing of olfactory information (Kay, 2015), converging evidences indicate that RROs occurs in a variety of higher-order cortical areas including the medial prefrontal cortex (mPFC, Biskamp et al., 2017, Nguyen Chi et al., 2016, Lockmann et al., 2016, Ito et al., 2014, Zhong et al., 2017, Karalis and Sirota, 2018, Moberly et al., 2018, Kőszeghy et al., 2018, Bagur et al., 2021). These results suggest that rhythmic breathing might affect cognitive functions beyond the processing of smells (Heck et al., 2019). Behavioural studies on human participants indeed demonstrated that nasal respiration supports memory encoding and recall (Zelano et al., 2016, Nakamura et al., 2018, Arshamian et al., 2018), but how respiration affects information processing and fundamental circuit operations in higher-order neocortex has remained largely unexplored on the mechanistic level.

Neuronal assemblies are thought to comprise the building blocks of cognitive function (Buzsáki, 2010, Papadimitriou et al., 2020, El-Gaby et al., 2021). Assemblies are composed of co-active neurons which transiently and consistently fire together, and are thought to convey information to downstream reader neurons by effective synaptic transmission due to their synchronized activity (Buzsáki, 2010). The recurrent nature of connections among cortical pyramidal cells and strengthening of connections of coactive neurons are thought to provide the structural and functional grounds for the emergence of assemblies (Harris, 2005, Palm et al., 2014). One way in which RROs could impact cortical information processing is to directly modulate the activity of assemblies. Focusing on the medial prefrontal cortex (mPFC), a highly associative brain area providing top-down control to cortex (Le Merre et al., 2021), we tested this hypothesis in awake mice. We find that assembly patterns emerge during spontaneous behaviour in the mPFC, and that these patterns are entrained by ongoing RROs. Assembly patterns preferentially activate during the descending phase of the RRO, when cortical excitation is maximized. Moreover, we provide evidence that the differential recruitment of putative GABAergic interneurons by assemblies during the descending phase of RRO supports the temporal segregation of assembly patterns. These results thus suggest that rhythmic breathing affects cognitive function at a fundamental level by defining time windows of preferred assembly activation.

## Results

### Prefrontal assemblies are entrained by respiration-related oscillations

The cortical local field potential (LFP) is characterized by prominent RROs, which peak in the 1-5 Hz frequency band during immobility (Biskamp et al., 2017, Zhong et al., 2017, Karalis and Sirota, 2018). We confirmed this finding in head-fixed mice, in which we simultaneously recorded LFP signals from the olfactory epithelium (LFP_olf_) and the mPFC (Karalis and Sirota, 2018, Fig. 1A,B). During immmobility, the mPFC LFP showed a spectral peak at ~1-5 Hz, which coincided with high spectral power of the LFP_olf_ (Fig. 1C,D). Furthermore, we found both signals to be coherent in the 1-5 Hz band (Fig. 1E). 1-5 Hz LFP power (Fig. 1C,F) and coherence with respiration moreover showed a dorso-ventral increase, consistent with a previous report on RROs in the mPFC (Karalis and Sirota, 2018, Fig. 1E,H). Our data thus support recent accounts that 1-5 Hz oscillatory activity in the mPFC reflects primarily a respiration-related rhythm. During movement, power spectra of both cortical LFP and respiration peaked at ~7-10 Hz (Supplementary Fig. 1). Given the spectral overlap to theta oscillations in mice (~7 Hz), it was less clear to what extend 7-10 Hz LFP oscillations were driven by respiration during movement. We thus focused on immobile states to assess the potential impact of 1-5 Hz RROs on neuronal assemblies.

**Fig. 1:**
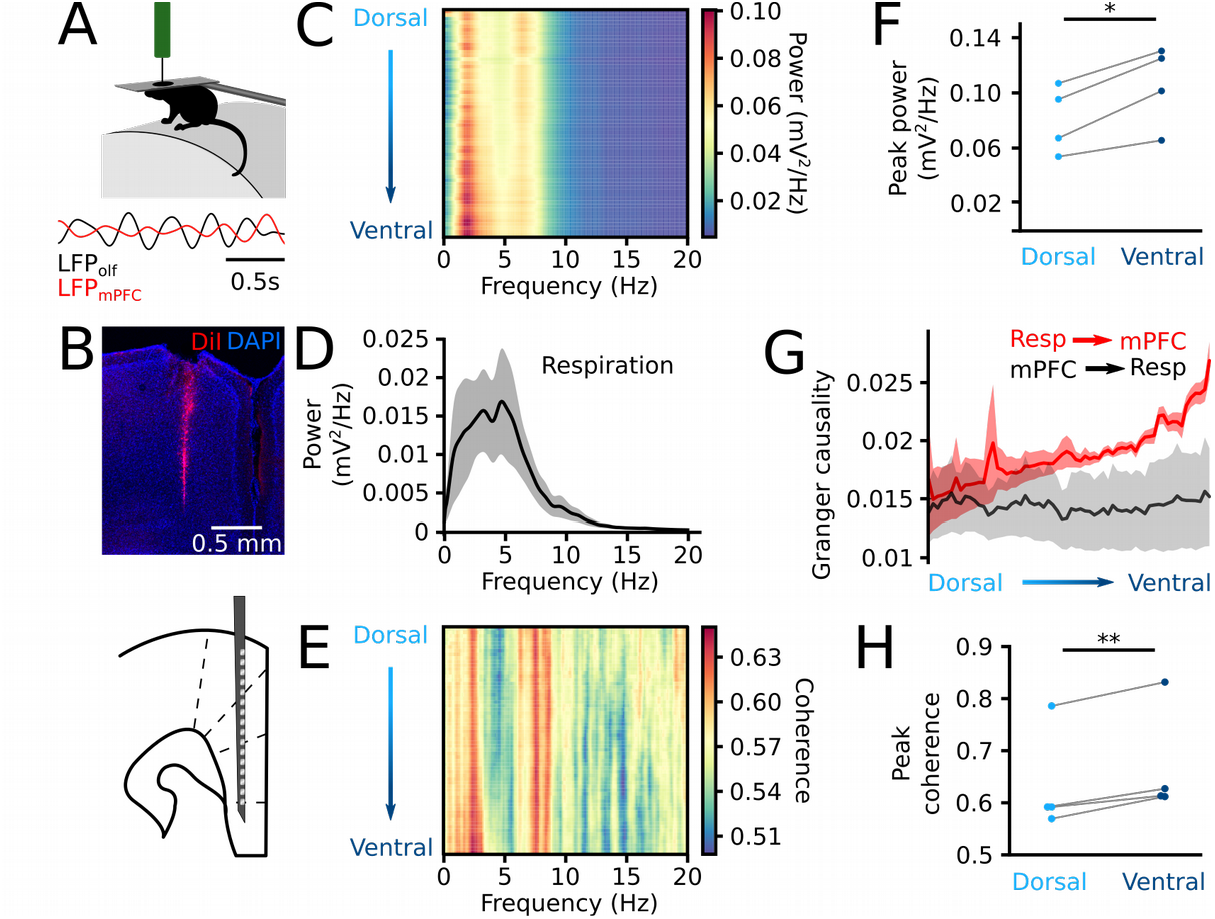
The mPFC LFP is entrained by respiration during immobility. A: Head-fixed mice were recorded during spontaneous immobility, bottom: example traces of the olfactory epithelium LFP (black) and the LFP in the mPFC (red) band pass filtered 1-4 Hz during immobility. B: Top, coronal section of the mPFC showing the shank of the silicon probe. Bottom: schematic of the recording configuration in the mPFC. C: Mean power spectral density of the mPFC along the dorso-ventral axis. D: Power spectral density of the olfactory epithelium LFP (shaded area: sem). E: Mean coherence between the mPFC and the respiration along the dorso-ventral axis. F: Amplitude of the peak power for the most dorsal and ventral recording sites. P=0.014. G: Depth profile of the Granger causality of the mPFC by the respiration (red) and of the respiration by the mPFC (black). H: Peak coherence between the respiration and the mPFC LFP for the most dorsal and ventral recording sites. n=4 mice, p=0.007. * p<0.05,** p<0.01, paired *t*-tests.

Motivated by previous reports of spontaneously occurring cell assemblies in neocortex and hippocampus (Peyrache et al., 2009, Miller et al., 2014, El-Gaby et al., 2021), we screened for assembly patterns in a dataset of single unit recordings from the mPFC of head-fixed, awake mice navigating in a virtual arena. In this paradigm, the animals showed periods of voluntary locomotion intermingled with extended epochs of immobility (proportion immobility: 0.40 ± 0.03, n=13 mice). We identified neuronal assembly activations from the occurrence of co-firing of neurons exceeding random coactivation (25 ms bin width, 60 ± 4 neurons per session, Fig. 2A, Supplementary Fig 2, Lopes-dos-Santos et al., 2013). This approach reliably extracted cell assemblies in simulated data (Supplementary Fig. 2) and identified on average one assembly pattern per 6.8 ± 0.2 neurons in the mPFC data set (25 sessions from 13 mice, 1494 pyramidal cells in total), similar to results from the hippocampus (El-Gaby et al., 2021). Assembly patterns were dominated by few neurons with large weights, which displayed more strongly correlated spike trains than neurons with low weight (p=10^−131^, Fig. 2A, see Supplementary Fig. 3 for additional quantification of assembly parameters). Ongoing network activity in the mPFC is thus characterized by the emergence of spontaneously activating neuronal assemblies.

**Fig. 2:**
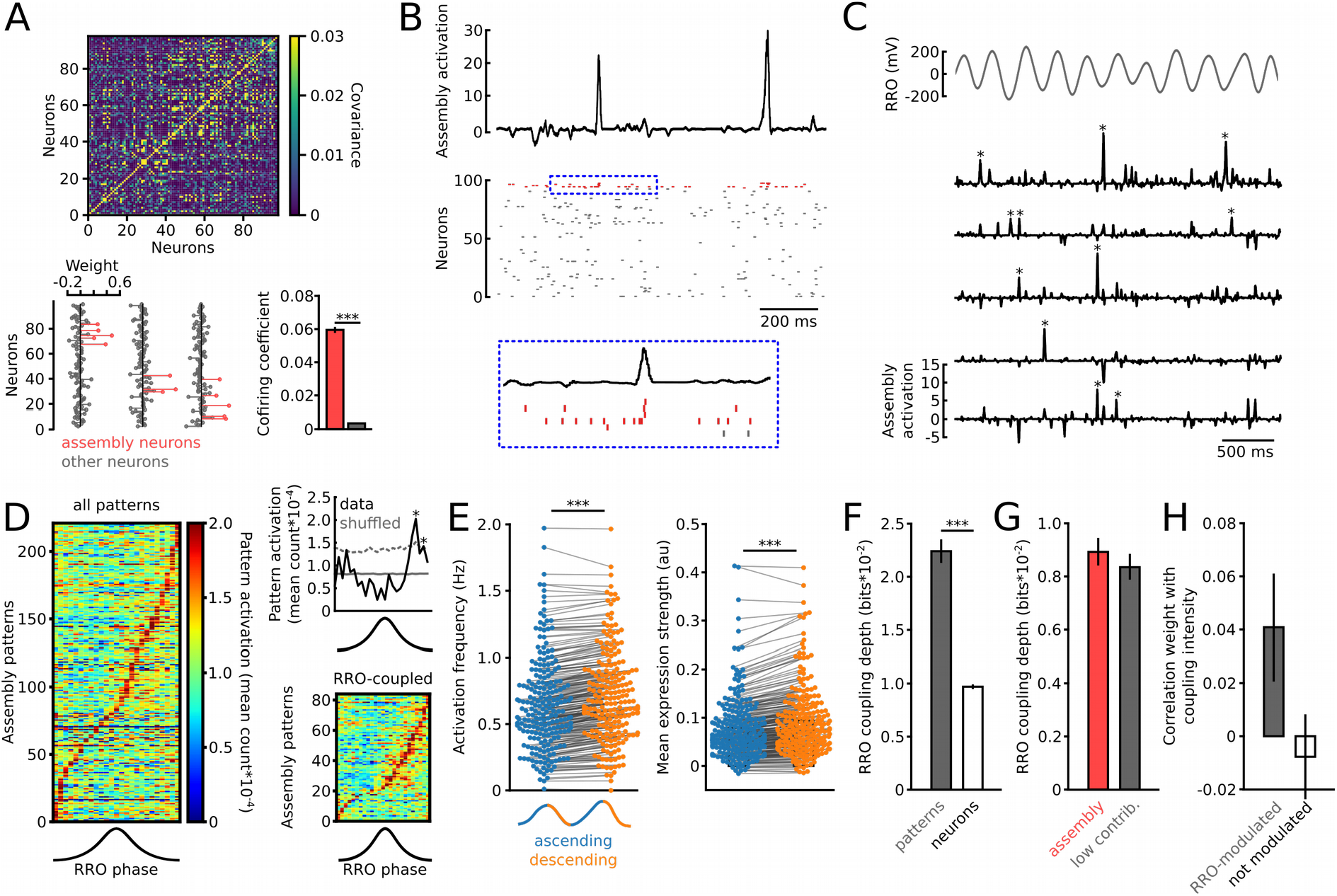
Cell assemblies preferentially activate during the descending phase of RROs. A: Assemblies were extracted from covariance matrices of binned spike trains. Top, the example covariance matrix of simultaneously recorded pyramidal cells in one session. Bottom, examples of three assembly weight vectors with assembly neurons labelled in red. Assembly neurons showed stronger cofiring than other neurons, confirming their joint assembly membership. n=674 assembly and 454206 other pairs, p=10^−131^. B: Time course of activation of the first assembly pattern shown in A along with the pyramidal cells giving rise to the pattern. The inset in the blue box shows one activation at higher resolution. C: Examples of assembly activations (asterisks) with simultaneously recorded RRO (1-5 Hz-filtered) during immobility. D: Average assembly activation frequency as a function of RRO phase revealed preferential occurrence of assemblies during the descending phase. Right: Example (top) and summary (bottom) of significantly RRO-entrained assemblies. E: Average pattern frequency (left) and expression strength (right) during the ascending and descending phase. n=221, p=4.8*10^−11^ and p=6.0*10^−6^ paired *t-*tests. F: RRO-coupling strenghts of assemblies and all individual pyramidal neurons. n=221 and 1145, p=6*10^−20^. G: Assembly neurons and cells with low contribution to a pattern showed similar RRO-coupling depth. n=142 and 143 patterns, p=0.648. H: Similar correlation between weight in an assembly pattern and RRO coupling intensity for RRO-modulated (n=84) and non-modulated patterns (n=137). p=0.063. *** p<0.001, Welch’s tests. Data are shown as mean ± sem.

To quantify the expression of assembly patterns with high temporal resolution, we extracted for each pattern the time course of activation by projecting the weight vectors on smoothed spike trains of all simultaneously recorded pyramidal neurons (Fig. 2B,C, van de Ven et al., 2016). We extracted all RRO cycles during which a given pattern activated, and quantified the number of assembly activations as a function of RRO phase. Importantly, since the duration of each phase bin is taken into account, this analysis is robust against waveform asymmetries of the RRO. Comparison against randomly shuffled onset times for each pattern revealed that 38 % of identified assembly patterns were significantly entrained by the ongoing RRO (84 out of 221 patterns, n=13 mice, Fig. 2D). The majority of patterns activated during the descending phase of RRO, thus coinciding with excitation of the circuitry during negative LFP deflections (Fig. 2D). This finding was robust against different threshold values for the detection of active assemblies (Supplementary Fig. 4). Considering all 221 patterns, we detected a significantly higher activation frequency and stronger average expression strength during the descending compared to the ascending phase (p=4.8*10^−11^ and p=6.0*10^−6^, Fig. 2E). These data thus demonstrate that RRO defines time windows of preferred activity for neuronal ensembles.

### RRO entrainment of assemblies emerges despite variable coupling of contributing neurons

We next asked whether the entrainment of assembly patterns by rhythmic breathing is a reflection of the functional grouping of highly RRO-coupled neurons into assemblies, or whether it is an emergent property that is independent from the RRO-coupling of the contributing neurons. We found evidence for the latter: First, the mean coupling strength of patterns was higher than that of individual neurons (p=6*10^−20^, Fig. 2F). Second, the average RRO coupling intensity of neurons with high contribution to assembly patterns did not differ from low-contributing neurons (p=0.648, Fig. 2G), indicating that coactivity of pyramidal cells with varying RRO coupling depth underlies RRO-paced assemblies. Third, correlation of a neuron’s weight in the assembly with the RRO coupling intensity of that neuron was generally low and showed no significant difference for RRO-entrained and non-entrained patterns (p=0.063, Fig. 2H). Thus, it is the transient coactivation of assemblies which is entrained by respiration, independently of the coupling of the individual neurons forming them.

### RRO-paced interneuron activity supports sparse assembly activations

Given that assemblies activate more often during the descending phase of RR, we next asked whether this results in enhanced assembly overlap due to an increase in co-occurence by chance. We quantified the coactivation of any two simultaneously recorded patterns within a time window of ±10 ms, which is within the integration time of cortical neurons (Koch et al., 1996). Despite higher assembly frequency (Fig. 2E), we observed reduced coactivation during the descending compared to the ascending phase (Fig. 3A, n=931 pairs of patterns, p=0.004). These data suggest that active mechanisms contribute to keeping assembly activations apart from each other during descending RRO.

**Fig. 3:**
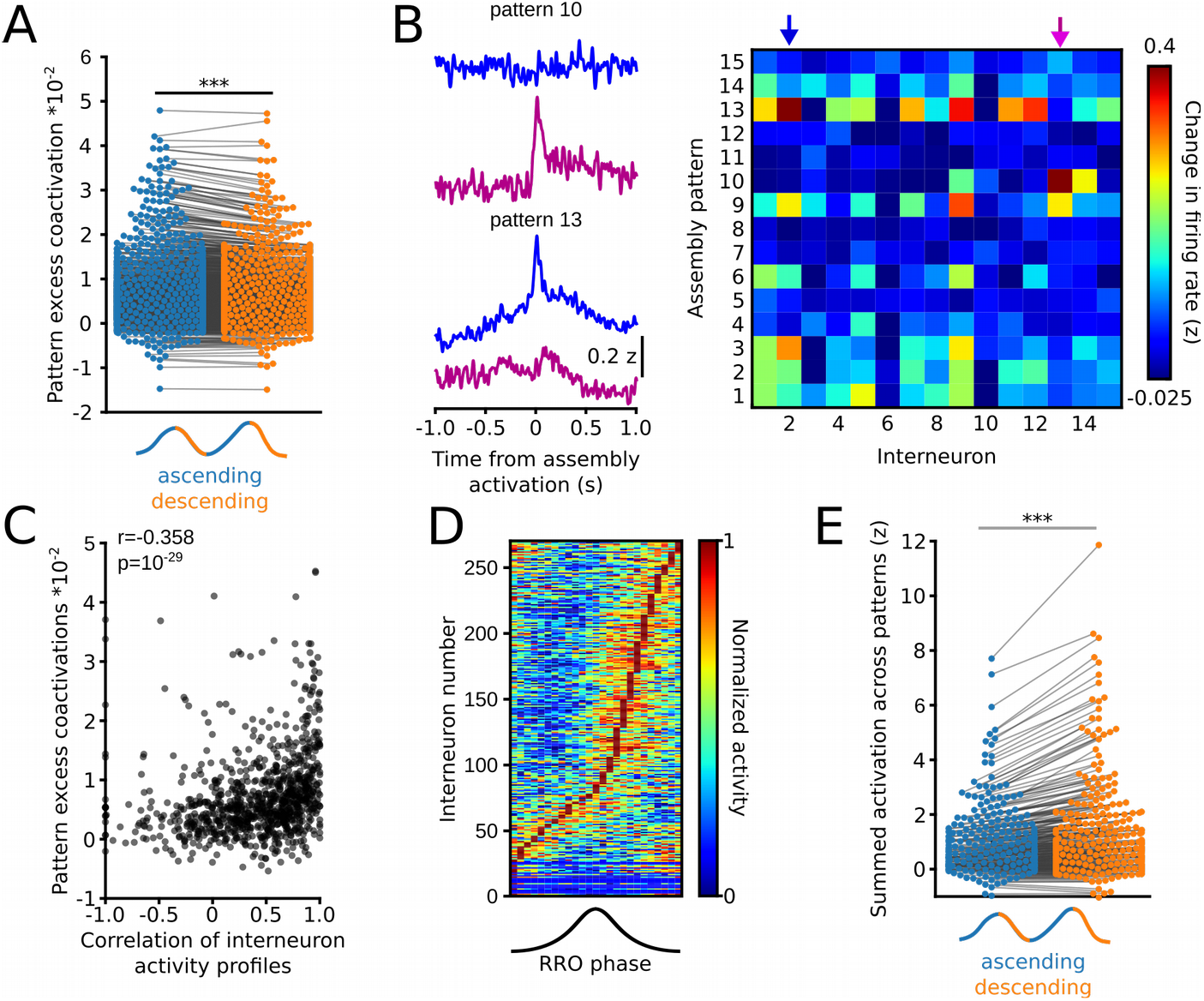
RRO phase-specific alignment of interneurons with assemblies. A: Quantification of coactivation of assembly patterns within ±10 ms revealed reduced coactivation frequency during the descending phase of ongoing RRO. n=931 pairs, p=0.004. B: Interneurons show differential activation profiles in relation to patterns. Left: example normalized firing rate of two interneurons (purple and blue) relative to the onset of two different assembly patterns (pattern 10 and 13). Right: Interneuron profile matrix summarizing the normalized firing change of all interneurons in the recording in response to the onset of all patterns in one session. Arrows indicate the interneurons shown on the left. C: Correlation of pattern coactivation strength and similarity of the interneuron profile of the same patterns. n=13 mice. D: Interneurons align their firing to the descending phase of RRO. E: Interneuron recruitment by assemblies is enhanced during descending RRO. The graph shows the summed normalized activation of each interneuron to all patterns. n=270 interneurons, p=6*10^−11^. Paired *t*-tests.

Previous work showed that GABAergic interneurons associate their activity with individual assembly patterns (Dupret et al., 2013). Feedback inhibition by GABAergic cells that are recruited by some but not other assemblies might thus provide a mechanism to maintain a sparse assembly activation profile (Buzsáki, 2010). For this to be true, interneurons should be differentially recruited by individual assembly patterns, fire when assemblies activate, and be more strongly aligned with assemblies during the descending phase of RRO. To test these predictions, we analyzed electrophysiolgically identified interneurons that were recorded simultaneously with the pyramidal cell population (n=270 putative interneurons). We found that interneurons showed diverse activity profiles (i.e. firing change in relation to the onset of each assembly pattern, Fig. 3B). The similarity in interneuron profiles between two patterns correlated positively with the coactivation strength of the same pair of patterns (Fig. 3C, Spearman’s *r*=0.358, p=3*10^−29^), indicating that strongly coactivating patterns share similar interneuron profiles. Moreover, similar to assembly patterns, interneurons discharged more during the descending phase of RRO (Fig. 3D). Finally, interneurons showed stronger coactivation with the assembly patterns during the descending than the ascending phase (Fig. 3E, n=270 interneurons, p=4*10^−11^). Jointly these data suggest that the pattern-specific coactivation of interneurons with individual assembly patterns provides a mechanism to support the segregation of assemblies during the descending phase of RRO.

### Phase-specific recruitment of interneurons by assembly neurons

Finally, we asked which mechanisms might mediate the enhanced assembly-recruitment of interneurons during descending RRO. One possibility would be that interneurons become more responsive to local glutamatergic drive from assembly neurons. To directly test this hypothesis, we analyzed putative excitatory synaptic connections from pyramidal cells onto interneurons using spike train cross-correlation (English et al., 2017). In total, we detected 234 connections (Fig. 4A, n=13 mice, 14842 connections tested). During the descending phase or RRO, spike transmission probability was significantly increased compared to the ascending phase (Fig. 4B, n=204 synaptic interactions, p=0.009). To directly compare assembly and non-assembly neurons, we separated the data set in connections from pyramidal neurons with high weight in at least one pattern in the recording (assembly connections, ~39% of connections) or low weight (non-assembly connections, 61% of connections). Both types of connections did not differ in their overall spike transmission (Fig. 4C, p=0.532), connection probability, or convergence (Supplementary Fig. 5). However, while non-assembly connections showed indistinguishable spike transmission when analysed separately for the ascending and descending phase of RRO (Fig. 4D, n=122, p=0.222), assembly connections displayed stronger spike transmission probability during the descending phase (Fig. 4D, n=82, p=0.003). These data jointly suggest that RROs support sparse assembly activity during the descending phase of each cycle by defining time windows of enhanced responsiveness of the local interneuron population to excitatory drive from assembly neurons.

**Fig. 4:**
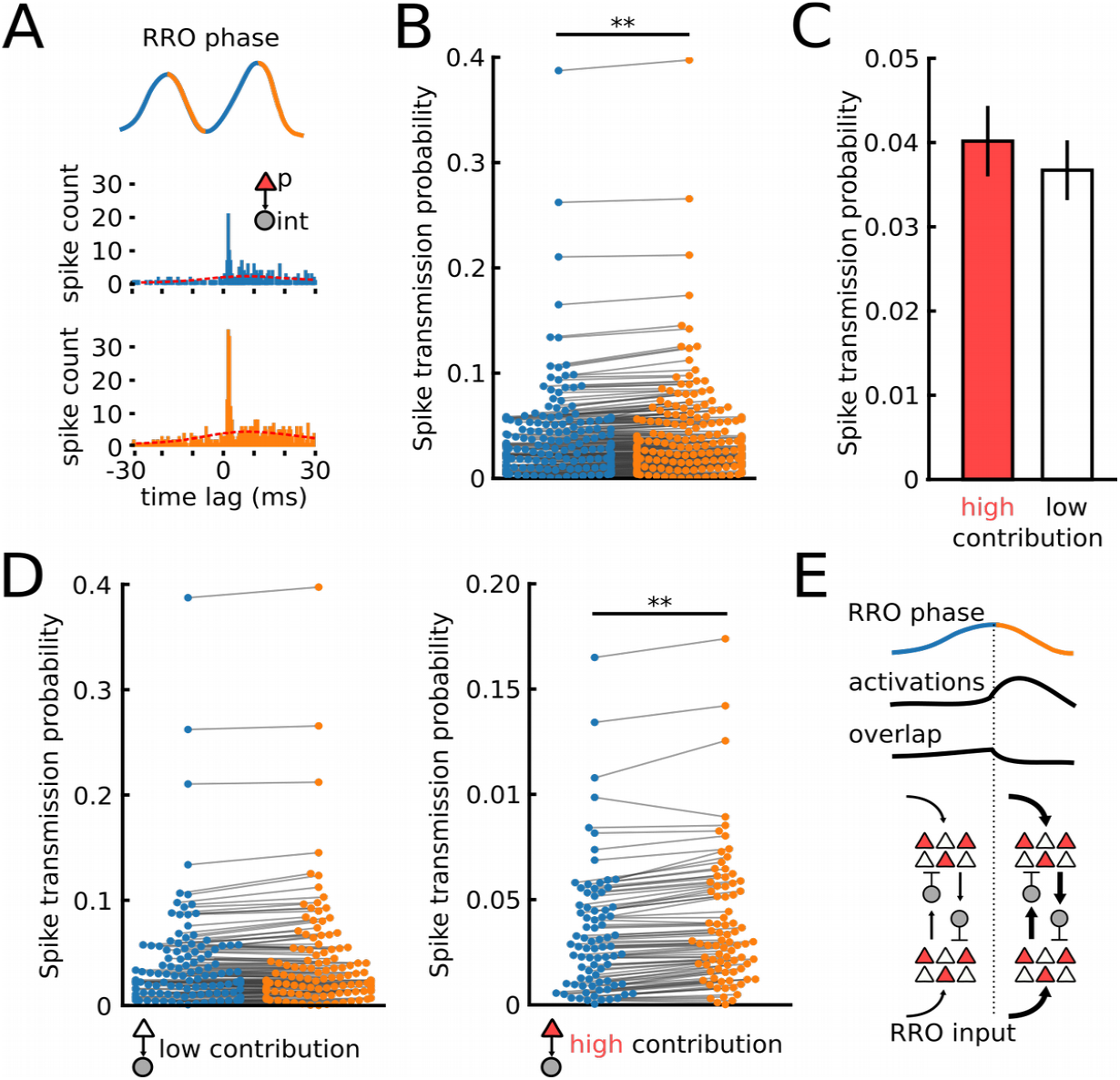
RRO phase-specific recruitment of interneurons by assembly neurons. A. Example of spike transmission at a pyramidal cell (p)-interneuron (int) connection during the ascending (blue) and descending phase of RRO (orange). Red line indicates the slowly comodulated baseline. B: Summary of connection strength during ascending and descending RRP phase. Spike transmission was significantly enhanced during the descending phase. n=204 connections, p=0.009. C: Unaltered overall spike transmission probability at assembly neuron-interneuron (high contribution, n=91) and non-assembly-interneuron connections (low contribution, n=143, p=0.532, Welch’s test). D: Spike transmission for non-assembly neuron-to-interneuron connections did not depend on RRO phase (left, n=122, p=0.222) while connections from assembly neurons displayed stronger transmission during descending RRO (right, n=82, p=0.0027). E: Schematic of the proposed function of RRO: Assembly activations are favoured during the descending phase, while assembly overlap is minimized due to the enhanced recruitment of interneurons. **p<0.01. Data in C are shown as mean ± sem. Paired *t*-tests unless indicated otherwise.

## Discussion

We found that spontaneously occurring assemblies in the mPFC align their activation with the descending phase of ongoing RRO. We provide evidence that this effect is explained by an emergent property of the circuit rather than the simple combination of RRO-coupled neurons into assemblies. This implies that RROs might have a stronger pacemaking effect on neuronal circuits than previously assumed on the basis of recordings from individual neurons.

A technical limitation when studying neuronal assemblies is the unequivocal identification of time points of assembly onset. Incomplete sampling of the local neuron population and thresholding the assembly expression time course, as done in our study and others (van de Ven et al., 2016, El-Gaby et al., 2021), might underestimate the real number of active assemblies at any given time point. However, our key finding that RROs define time windows of preferred assembly activation did hold for different activation thresholds, suggesting that the RRO modulation does not depend on the parameter selection but rather represents a fundamental property of the behaviour of cortical assemblies. It should be further noted that our method of assembly detection does not take into account the temporal structure of the neuronal activity in the assemblies, but solely detects whether or not neurons show significant coactivation. Neuronal assemblies have, however, also been defined based on the temporal alignment of spikes (i.e. neuronal sequences) in both the hippocampus (Chenani et al., 2019) and neocortex (Carrillo-Reid et al., 2015, Luczak et al., 2007, Luczak et al., 2009). Future work will be required to test whether the principle of RR modulation applies to such neuronal sequences.

Our data add to the notion that cell assemblies exist in the absence of specific stimuli, arguing in favor of pre-existing network structure suitable to integrate new information using a pool of readily available network motifs (Miller et al., 2014, Almeida-Filho et al., 2014, Carrillo-Reid et al., 2015, Hamm et al., 2017, Dejean et al., 2016). While assembly activity was higher during the descending RRO phase, the overlap between individual assemblies was reduced during that time (Fig. 3A). These data imply that the co-activation of assemblies is actively suppressed during the descending phase of RR, as the higher frequency of assembly activation would make random coactivity more likely. The simultaneous recruitment of putative GABAergic interneurons, which happens distinctly for different assembly patterns, provides a potential mechanism how individual assemblies might separate their activation from each other (Buzsáki, 2010). These data are in agreement with results from the hippocampus showing that GABAergic neurons differentially align their firing with cell assemblies representing distinct places during spatial learning (Dupret et al., 2013). In line with this hypothesis, interneurons were recruited more strongly to assemblies during the descending phase of RRO (Fig. 3E). Importantly, although occurring on the basis of higher interneurons rates during descending RRO, the enhanced interneuron recruitment was not merely an effect of higher interneuron firing since our measure of assembly-aligned recruitment takes into account the local rate before assembly onset. The association of interneuron firing with assemblies could be caused by an enhanced excitability state due to impinging respiration-driven excitation linked with negative LFP deflections in combination with short-term plasticity processes. To lines of evidence argue in favor of this hypothesis: First, using a cross-correlation-based estimation of spike transmission probability, we show that interneurons are particularly receptive to incoming excitatory signals from assembly neurons (Fig. 4D). Presynaptic cooperativity could synergistically impact spike transmission at assembly neuron-interneuron connections: In the hippocampus, synchronized presynaptic activity leads to enhanced spike transmission (English et al., 2017). Such synchronization would be expected for assembly neurons and might thus contribute to higher spike transmission during the descending phase of RROs. Second, interneurons have been shown to be particularly receptive to respiration-driven input. They are consistently found to be more likely to be phase-coupled to RRO than principal cells (Karalis and Sirota, 2018, Biskamp et al., 2017). Furthermore, whole-cell recordings from pyramidal cells in the parietal cortex revealed subthreshold respiration-synchronous membrane potential oscillations likely reflecting GABAergic synaptic currents (Jung et al., 2019). These findings imply that the main effect of respiration-driven inputs to neocortical circuits is mediated through GABAergic neurons.

Our results add to the increasing recognition of internally generated bodily influences as modulators of brain activity and cognitive functions, including drive from respiration, heart rate and gastrointestinal rhythms (Heck et al., 2019, Azzalini et al., 2019). RROs have been directly observed in various areas of the neocortex, hippocampus, thalamus, and amygdala (Zhong et al., 2017, Ito et al., 2014, Biskamp et al., 2017, Nguyen Chi et al., 2016, Lockmann et al., 2016, Moberly et al., 2018, Jung et al., 2019, Bagur et al., 2021) and are thought to impact cortical circuits through the entrainment of brain oscillations involved in cognitive functions, including theta (Zelano et al., 2016), gamma (Zhong et al., 2017, Biskamp et al., 2017) and sharp-wave/ripple oscillations (Liu et al., 2017). Based on our results we propose that the role of RROs extends to the building blocks of cortical computations, the assemblies. The synchronization of assemblies to RROs might provide an effective sender-reader interaction such that the impact of synchronized activity from upstream can be efficiently interpreted by downstream reader implementations across neocortex and subcortical structures (Buzsáki, 2010). Alternatively, pooling sparse assembly activations in the descending phase of RRO might provide a mechanism to facilitate spontaneous assembly reactivations during offline states, which has been argued to support memory persistence in the presence of synaptic turnover (Fauth and van Rossum, 2019).

## Acknowledgements

This work was supported by the German Research Foundation (grant SA 3609/1-1 to J.-F.S) and by institutional funding (Institute of Physiology I).

## Author contributions

S.F. and J.-F.S. performed experiments, analyzed data, and wrote the manuscript. J.-F.S. designed the study.

## Methods

### Mice

C57Bl6/J mice of both sexes were used in this study. The animals had free access to food and water and were maintained on a 12 dark/light cycle. Mice were 6 to 13 weeks old. All experiments were performed in agreement with national legislation and were approved by the Regierungspräsidium Freiburg. We analyzed data from 10 mice that were recorded in the context of a previous study (Sauer and Bartos, 2021). 4 additional mice were implanted and recorded for this study.

### Surgical procedures

A stainless steel head plate was implanted on the skull under general anesthesia in isoflurane (induction: 3%, maintenance: 1-2%) using dental cement. In 4 mice, a 0.8 mm hole was drilled above the nasal cavity, and a silver wire insulated up to ~0.5 mm from the extremity was inserted between the olfactory epithelium and the bone, and cemented in place. The animals were allowed to recover from head plate implantation for at least three days. Buprenorphin (0.1 mg/kg body weight) and Carprofen (5 mg/kg body weight) were injected subcutaneously before the surgery for pain relief. Once the animals were habituated to head-fixation (see below), a craniotomy was performed over both mPFCs (1.9 mm anterior, 0.4 mm lateral of bregma) under isoflurane anesthesia. The craniotomy was then covered with a drop of phosphate-buffered saline (PBS) and sealed of with QuikCast elastomer until the recordings took place. An additional injection of Carprofen was given for analgesia prior to craniotomy.

### Single-unit recording in the virtual reality

The mice were habituated to running on a circular track in a virtual reality. For head fixation, the mice were briefly anesthetized in isoflurane (3% in O_2_). During habituation and recording, the mice ran on a circular styrofoam wheel. First, animals were habituated to head-fixation for at least three days without the virtual reality turned on. Then, they were daily exposed to the virtual reality (circular track, length 2-3 m, with visual cues placed outside the arena). The virtual reality was constructed with open-source 3D rendering software Blender (Schmidt-Hieber and Häusser, 2013) and was projected on five computer screens surrounding the head-fixation setup.

Recordings were performed 1-3 days after the craniotomy using H3 single-shank silicon probes (64 recording sites spaced 20 μm apart, total shank length 1275 μm, Cambridge Neurotech). The probe was coated with a fluorescent marker (DiI or DiO) and was slowly (~2-5 μm/s) lowered to the mPFC (1600-1900 μm below brain surface). The probe was left in place before the recordings for 10-15 min. Wide-band signals were recorded with a 64-channel amplifier (Intan Technologies) using OpenEphys GUI software (30 kHz sampling frequency). Movement of the animal was assessed by recording the motion of the running wheel as a pulse-width modulated signal. After the recording, the probe was slowly retracted and the craniotomy sealed off with QuickCast elastomer. With each mouse, 1-3 recording sessions were performed (1 session per day).

### Histology

After recording, the animals were deeply anesthetized with ketamin/xylazine (i.p. injection) and transcardially perfused with ~20 ml phosphate-buffered saline followed by ~30 ml of 4% paraformaldehyde. 100 μm-thick coronal sections of the mPFC were cut after post-fixation in fixative overnight at 4°. The slices were washed in PBS and stained with 4′,6-diamidino-2-phenylindole. A laser scanning microscope (LSM 710 or 900, Zeiss) was used to visualize the location of the silicon probe. Recording locations ranged from layer 2 to 6, spanning from the accessory motor cortex to the medial orbital area.

### Single-unit isolation

Single units were extracted from bandpass-filtered data (0.3-3 kHz) using MountainSort (Chung et al., 2017). Putative single-unit clusters with high isolation index (>0.90) and low noise overlap (<0.1) were kept for manual curation, during which only clusters with a clear refractory period were kept. In case of two clusters with similar waveforms, cross-correlation was used to assess whether clusters had to be merged. Isolated units were separated in putative excitatory and inhibitory neurons based on trough-to-peak duration and asymmetry index as described before (Sirota et al., 2008).

### Analysis of respiration-local field potential coupling

Olfactory epithelium and mPFC LFP power spectral density and cross spectral density were computed using Welch’s average periodogram method. For the power, the signals were divided in Hann windows of 2 s length with no overlap and padded by a factor 10, and the obtained power spectral density was then averaged across windows. The coherence was computed on windows of 4 s as the normalized cross spectral density of 2 s Hann windows with no overlap and then averaged. The Granger causality was defined as the variance of the residual from a linear auto-regressive model fitted on a 2 s window of the mPFC LFP or the LFP_olf_ divided by the residual of a vector auto-regressive model including the LFP_olf_ or the mPFC LFP. Both auto-regressive and vector auto-regressive models (from the *statsmodels* python package) had a fixed lag of 1, and were computed on signals filtered below 50 Hz and downsampled to 100 Hz.

### Detection of assembly patterns

Assembly extraction was performed using principal- and independent component analysis following a published procedure using simultaneously recorded pyramidal cells (van de Ven et al., 2016). The extraction was done on the entire recording duration, including movement and immobility, while the analysis of assembly activations in relation to ongoing RROs was performed during immobility. We used a total of 13 mice with sufficient immobility duration for this analysis.

The spike trains of *n* pyramidal neurons were binned in *B* 25 ms bins and normalized by z-scoring to avoid bias by highly active neurons. To detect the number of assembly patterns in a recording, principal component analysis was applied to the binned spike train matrix. We next used the Marčenko-Pastur law to extract the number of significant assembly patterns (Marčenko and Pastur, 1967, Lopes-dos-Santos et al., 2013). The Marčenko-Pastur law indicates that a correlation matrix constructed from independent random variables yields eigenvalues below a critical value *c* given as

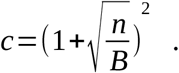

If neurons fire correlated with each other (as it would be the case for assemblies), eigenvalues above the critical limit will exist. The number of eigenvalues exceeding the theoretical limit thus indicates the number of assembly patterns (Lopes-dos-Santos et al., 2013). Independent component analysis was then used to extract activity patterns that are as independent from each other as possible. Using the fastICA algorithm of *scikit.learn*, we extracted the number of independent components given by the eigenvalues above *c.* The resulting components represent the weight vectors of each assembly pattern. Note that the orientation of independent components is arbitrary, so each vector was oriented to have the largest deflection in positive direction and was further scaled to unit length. Assembly neurons were defined as those cells with a weight exceeding 2x the standard deviation of the pattern vector (van de Ven et al., 2016). Sparsity of assembly patterns (i.e. to what extent assemblies were dominated by high weights of few neurons) was quantified as

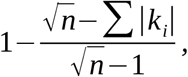

where n denotes the length of the weight vector, and *k_i_* is the weight of neuron *i* in pattern *k.*

### Reconstruction of assembly activations over time

To obtain the assembly activation time course *T* for all *k* patterns at high resolution, the weight vectors corresponding to the assemblies were projected on smoothed spike trains *z* of all neurons:

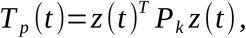

where T denotes the transpose operator and *P_k_* gives the outer product of the k^th^ weight vector. The spike train matrix *z* was constructed by convolving each neuron’s spike train with a Gaussian kernel (standard deviation 7.2 ms). This procedure resulted in smooth time courses of pattern activation. We set a threshold of 5 to detect assembly activations, unless indicated otherwise (van de Ven et al., 2016).

### Assembly detection in simulated data

A simulated binned spike train matrix *B_sim_* consisting of 70 neurons and 1000 bins was constructed as 70 Poisson neurons using the *numpy.random.poisson* function (with lam=1). Assemblies were modeled as a group of neurons with joint elevation in spike rate in 50 randomly chosen bins. The spike rate increase was modeled by randomly drawing a spike value ranging between 0 and 6 for each of the assembly neurons. This way, the identity of the assembly neurons was known a priori, while the time points of activation were not. The assembly extraction procedure was applied as described above, except that the reconstruction of the time course of the assemblies was done directly on *B_sim_* rather than convolved spike trains.

### Single-neuron analysis

The cofiring coefficient was calculated using Pearson’s correlation coefficient from binned spike trains (25 ms bin width) in a round-robin fashion separately for assembly and non-assembly neurons. To assess the spatial extent of assemblies, we measured for each pattern the average distance between all assembly neurons and a matching number of randomly drawn neurons. The position of the neuron was defined by the electrode with largest negative voltage deflection. RRO-coupling of units was quantified using the Kullback-Leibler distance (see below). Only cells with at least 200 spikes during the immobility epochs were considered for this analysis. To compare RRO-coupling for assembly and non-assembly neurons, the coupling value of assembly neurons for each pattern was compared with a matching number of low-contributing neurons (i.e. with the lowest weights in the pattern vector). The association of interneurons with assembly patterns was tested by first z-scoring the convolved interneuron spike trains. Then, the mean firing rate change during a 30 ms window following assembly onset relative to preceding baseline (833 ms long, ending 166 ms before assembly onset) was calculated for each pattern and interneuron.

### Assembly analysis

Assembly-RRO coupling was assessed by extracting first the times of assembly activations by threshold-crossing. After an onset was detected, no further activations could be scored for 50 ms to avoid double-detection. Then, for each activation we determined the instantaneous phase of the ongoing 1-5 Hz filtered and Hilbert-transformed RRO, and quantified the mean activation as a function of RRO phase bins (25 bins). This coupling measure thus carries the unit “mean activations/14.4°”. Coupling strength was expressed with the Kullback-Leibler distance *K* between the actual phase distribution *P* and a uniform distribution *U* with the same mean:

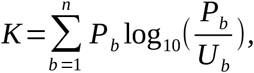

where *n* denotes the bin number. Significant coupling was tested by randomly shuffling the activation times (1000 iterations). Phase-coupling was considered significant when *K* exceeded the 99^th^ percentile of the random distribution. Kullback-Leibler distance was also used to obtain phase-coupling of single-units.

### Assembly coactivation and synaptic interactions

To detect assembly coactivations and putative pyramidal cell-interneuron connections, we used a cross-correlation based framework (English et al., 2017). For synaptic connections, we first determined the raw cross-correlation between two binned spike trains (0.4 ms bins) for neurons with more than 500 spikes using the filter_correlogram function of the *neuronpy.util.spiketrain* package. Criteria for a significant monosynaptic interaction were a peak in the monosynaptic time window (0.8-2.8 ms following the spike in the pyramidal cell) significantly exceeding the co-modulated baseline and the peak in anti-causal direction (i.e. interneuron-pyramidal cell, −2 to 0 ms). The baseline *b* was obtained by convolving the raw cross-correlogram with a partially hollowed Gaussian function (hollow fraction: 0.6, standard deviation: 10 ms). The Poisson distribution with continuity correction was used to estimate the probability of the observed magnitude of cross-correlation in the monosynaptic bins (*P_syn_*),

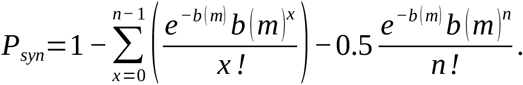

Similarly, we estimated the probability of the observed count in the monosynaptic bins of the cross-correlogram being larger than the count in anticausal direction (c_anticausal_) using the Poisson distribution with continuity correction,

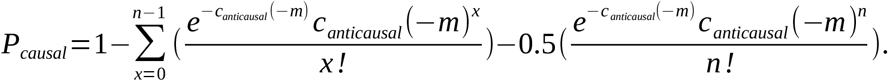

Following optogenetic ground truth data obtained in the hippocampus, a pair was marked as connected if P_syn_<0.001 and P_causal_<0.0026 (English et al., 2017). Spike transmission probability was defined as the spiking in the monosynaptic window exceeding *b* normalized by the number of presynaptic spikes. For all significantly connected pairs, we additionally extracted spike transmission probability separately for the ascending and descending phases of RRO, which were defined from 1-5 Hz filtered and Hilbert-transformed raw LFP traces. Only connections with positive spike transmission during both ascending and descending phase were considered for this analysis. Convergence was assessed by taking the number of convergent connections divided by the number of total connections of the session. This analysis was only applied to sessions with at least 3 connections (12 sessions from 10 mice). For assembly coactivations, the coactivation strength was determined by summing the values exceeding *b* in the −10 to + 10 ms time window for all pairs of patterns.

### Statistical analysis

Unpaired comparisons were done with two-sided Welch’s tests, which is robust against deviation from normal distribution at large sample sizes (Stonehouse and Forrester, 1998). For small group sizes <15, an unpaired two-sided *t*-test was used. Correlations were assessed with Spearman’s correlation coefficient. Pairwise comparisons were done with a paired *t-*test. Data are presented as full data ranges or as mean ± sem where indicated. All analysis (except for initial spike sorting) were performed in Python2.7.

## References

Almeida-Filho DG, Lopes-dos-Santos V, Vasconcelos NAP, Miranda JGV, Tort ABL, Ribeiro S. 2014. An investigation of Hebbian phase sequences as assembly graphs. Front Neural Circuits 8. doi:10.3389/fncir.2014.00034

Arshamian A, Iravani B, Majid A, Lundström JN. 2018. Respiration Modulates Olfactory Memory Consolidation in Humans. J Neurosci 38:10286–10294. doi:10.1523/JNEUROSCI.3360-17.2018

Azzalini D, Rebollo I, Tallon-Baudry C. 2019. Visceral Signals Shape Brain Dynamics and Cognition. Trends in Cognitive Sciences 23:488–509. doi:10.1016/j.tics.2019.03.007

Bagur S, Lefort JM, Lacroix MM, de Lavilléon G, Herry C, Chouvaeff M, Billand C, Geoffroy H, Benchenane K. 2021. Breathing-driven prefrontal oscillations regulate maintenance of conditioned-fear evoked freezing independently of initiation. Nat Commun 12:2605. doi:10.1038/s41467-021-22798-6

Biskamp J, Bartos M, Sauer J-F. 2017. Organization of prefrontal network activity by respiration-related oscillations. Sci Rep 7:45508. doi:10.1038/srep45508

Buzsáki G. 2010. Neural syntax: cell assemblies, synapsembles, and readers. Neuron 68:362–385. doi:10.1016/j.neuron.2010.09.023

Carrillo-Reid L, Miller JK, Hamm JP, Jackson J, Yuste R. 2015. Endogenous Sequential Cortical Activity Evoked by Visual Stimuli. J Neurosci 35:8813–8828. doi:10.1523/JNEUROSCI.5214-14.2015

Chenani A, Sabariego M, Schlesiger MI, Leutgeb JK, Leutgeb S, Leibold C. 2019. Hippocampal CA1 replay becomes less prominent but more rigid without inputs from medial entorhinal cortex. Nature Communications 10:1341. doi:10.1038/s41467-019-09280-0

Chung JE, Magland JF, Barnett AH, Tolosa VM, Tooker AC, Lee KY, Shah KG, Felix SH, Frank LM, Greengard LF. 2017. A Fully Automated Approach to Spike Sorting. Neuron 95:1381–1394.e6. doi:10.1016/j.neuron.2017.08.030

Dejean C, Courtin J, Karalis N, Chaudun F, Wurtz H, Bienvenu TCM, Herry C. 2016. Prefrontal neuronal assemblies temporally control fear behaviour. Nature 535:420–424. doi:10.1038/nature18630

Dupret D, O’Neill J, Csicsvari J. 2013. Dynamic reconfiguration of hippocampal interneuron circuits during spatial learning. Neuron 78:166–180. doi:10.1016/j.neuron.2013.01.033

El-Gaby M, Reeve HM, Lopes-Dos-Santos V, Campo-Urriza N, Perestenko PV, Morley A, Strickland LAM, Lukács IP, Paulsen O, Dupret D. 2021. An emergent neural coactivity code for dynamic memory. Nat Neurosci 24:694–704. doi:10.1038/s41593-021-00820-w

English DF, McKenzie S, Evans T, Kim K, Yoon E, Buzsáki G. 2017. Pyramidal Cell-Interneuron Circuit Architecture and Dynamics in Hippocampal Networks. Neuron 96:505–520.e7. doi:10.1016/j.neuron.2017.09.033

Fauth MJ, van Rossum MC. 2019. Self-organized reactivation maintains and reinforces memories despite synaptic turnover. eLife 8:e43717. doi:10.7554/eLife.43717

Fontanini A, Bower JM. 2006. Slow-waves in the olfactory system: an olfactory perspective on cortical rhythms. Trends Neurosci 29:429–437. doi:10.1016/j.tins.2006.06.013

Grosmaitre X, Santarelli LC, Tan J, Luo M, Ma M. 2007. Dual functions of mammalian olfactory sensory neurons as odor detectors and mechanical sensors. Nat Neurosci 10:348–354. doi:10.1038/nn1856

Hamm JP, Peterka DS, Gogos JA, Yuste R. 2017. Altered Cortical Ensembles in Mouse Models of Schizophrenia. Neuron 94:153–167.e8. doi:10.1016/j.neuron.2017.03.019

Harris KD. 2005. Neural signatures of cell assembly organization. Nat Rev Neurosci 6:399–407. doi:10.1038/nrn1669

Heck DH, Kozma R, Kay LM. 2019. The rhythm of memory: how breathing shapes memory function. J Neurophysiol 122:563–571. doi:10.1152/jn.00200.2019

Ito J, Roy S, Liu Y, Cao Y, Fletcher M, Lu L, Boughter JD, Grün S, Heck DH. 2014. Whisker barrel cortex delta oscillations and gamma power in the awake mouse are linked to respiration. Nat Commun 5:3572. doi:10.1038/ncomms4572

Jung F, Yanovsky Y, Brankačk J, Tort AB, Draguhn A. 2019. Respiration competes with theta for modulating parietal cortex neurons. bioRxiv 707331. doi:10.1101/707331

Karalis N, Sirota A. 2018. Breathing coordinates limbic network dynamics underlying memory consolidation. bioRxiv 392530. doi:10.1101/392530

Kay LM. 2015. Olfactory system oscillations across phyla. Curr Opin Neurobiol 31:141–147. doi:10.1016/j.conb.2014.10.004

Koch C, Rapp M, Segev I. 1996. A brief history of time (constants). Cereb Cortex 6:93–101. doi:10.1093/cercor/6.2.93

Kőszeghy Á, Lasztóczi B, Forro T, Klausberger T. 2018. Spike-Timing of Orbitofrontal Neurons Is Synchronized With Breathing. Front Cell Neurosci 12:105. doi:10.3389/fncel.2018.00105

Le Merre P, Ährlund-Richter S, Carlén M. 2021. The mouse prefrontal cortex: Unity in diversity. Neuron. doi:10.1016/j.neuron.2021.03.035

Liu Y, McAfee SS, Heck DH. 2017. Hippocampal sharp-wave ripples in awake mice are entrained by respiration. Sci Rep 7:8950. doi:10.1038/s41598-017-09511-8

Lockmann ALV, Laplagne DA, Leão RN, Tort ABL. 2016. A Respiration-Coupled Rhythm in the Rat Hippocampus Independent of Theta and Slow Oscillations. J Neurosci 36:5338–5352. doi:10.1523/JNEUROSCI.3452-15.2016

Lopes-dos-Santos V, Ribeiro S, Tort ABL. 2013. Detecting cell assemblies in large neuronal populations. J Neurosci Methods 220:149–166. doi:10.1016/j.jneumeth.2013.04.010

Luczak A, Barthó P, Harris KD. 2009. Spontaneous events outline the realm of possible sensory responses in neocortical populations. Neuron 62:413–425. doi:10.1016/j.neuron.2009.03.014

Luczak A, Barthó P, Marguet SL, Buzsáki G, Harris KD. 2007. Sequential structure of neocortical spontaneous activity in vivo. PNAS 104:347–352. doi:10.1073/pnas.0605643104

Marčenko VA, Pastur LA. 1967. DISTRIBUTION OF EIGENVALUES FOR SOME SETS OF RANDOM MATRICES. Math USSR Sb 1:457. doi:10.1070/SM1967v001n04ABEH001994

Miller JK, Ayzenshtat I, Carrillo-Reid L, Yuste R. 2014. Visual stimuli recruit intrinsically generated cortical ensembles. PNAS 111:E4053–E4061. doi:10.1073/pnas.1406077111

Moberly AH, Schreck M, Bhattarai JP, Zweifel LS, Luo W, Ma M. 2018. Olfactory inputs modulate respiration-related rhythmic activity in the prefrontal cortex and freezing behavior. Nat Commun 9:1528. doi:10.1038/s41467-018-03988-1

Nakamura NH, Fukunaga M, Oku Y. 2018. Respiratory modulation of cognitive performance during the retrieval process. PLoS One 13:e0204021. doi:10.1371/journal.pone.0204021

Nguyen Chi V, Müller C, Wolfenstetter T, Yanovsky Y, Draguhn A, Tort ABL, Brankačk J. 2016. Hippocampal Respiration-Driven Rhythm Distinct from Theta Oscillations in Awake Mice. J Neurosci 36:162–177. doi:10.1523/JNEUROSCI.2848-15.2016

Palm G, Knoblauch A, Hauser F, Schüz A. 2014. Cell assemblies in the cerebral cortex. Biol Cybern 108:559–572. doi:10.1007/s00422-014-0596-4

Papadimitriou CH, Vempala SS, Mitropolsky D, Collins M, Maass W. 2020. Brain computation by assemblies of neurons. PNAS 117:14464–14472. doi:10.1073/pnas.2001893117

Peyrache A, Khamassi M, Benchenane K, Wiener SI, Battaglia FP. 2009. Replay of rule-learning related neural patterns in the prefrontal cortex during sleep. Nat Neurosci 12:919–926. doi:10.1038/nn.2337

Sauer J-F, Bartos M. 2021. Topographically organized representation of space and context in the medial prefrontal cortex. bioRxiv 2021.06.04.447085. doi:10.1101/2021.06.04.447085

Schmidt-Hieber C, Häusser M. 2013. Cellular mechanisms of spatial navigation in the medial entorhinal cortex. Nat Neurosci 16:325–331. doi:10.1038/nn.3340

Sirota A, Montgomery S, Fujisawa S, Isomura Y, Zugaro M, Buzsáki G. 2008. Entrainment of neocortical neurons and gamma oscillations by the hippocampal theta rhythm. Neuron 60:683–697. doi:10.1016/j.neuron.2008.09.014

Stonehouse JM, Forrester GJ. 1998. Robustness of the t and U tests under combined assumption violations. Journal of Applied Statistics 25:63–74. doi:10.1080/02664769823304

van de Ven GM, Trouche S, McNamara CG, Allen K, Dupret D. 2016. Hippocampal Offline Reactivation Consolidates Recently Formed Cell Assembly Patterns during Sharp Wave-Ripples. Neuron 92:968–974. doi:10.1016/j.neuron.2016.10.020

Zelano C, Jiang H, Zhou G, Arora N, Schuele S, Rosenow J, Gottfried JA. 2016. Nasal Respiration Entrains Human Limbic Oscillations and Modulates Cognitive Function. J Neurosci 36:12448–12467. doi:10.1523/JNEUROSCI.2586-16.2016

Zhong W, Ciatipis M, Wolfenstetter T, Jessberger J, Müller C, Ponsel S, Yanovsky Y, Brankačk J, Tort ABL, Draguhn A. 2017. Selective entrainment of gamma subbands by different slow network oscillations. Proc Natl Acad Sci USA 114:4519–4524. doi:10.1073/pnas.1617249114

