## Supplementary_materials for "Phase-specific pooling of sparse assembly activity by respiration-related brain oscillations"

**Supplementary data:**

Supplementary Figures 1-5

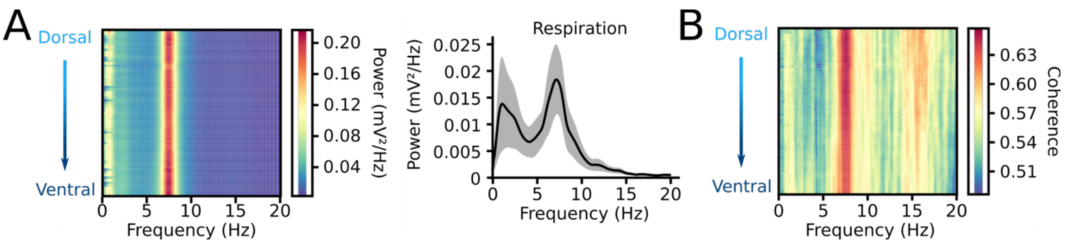


**Supplementary Fig. 1: LFP properties during movement.**

A: Average power spectral density of LFP power along the recording shank (left) and of respiration (right) during movement (>2 cm/s). n=4 mice. B: Coherence between prefrontal LFP and respiration during movement epochs. n=14 mice.

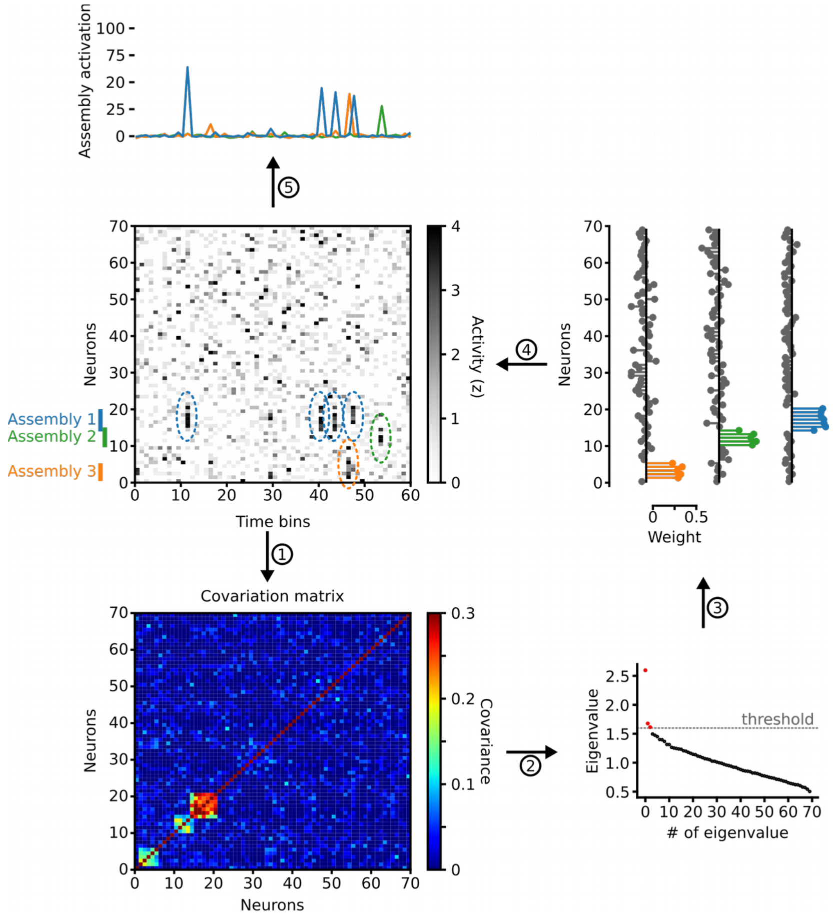


**Supplementary Fig. 2: Detection of assembly patterns.**

Three assemblies were inserted into a Poisson matrix of binned spike trains. Note that assembly 1 and 2 share a common neuron. Randomly timed assembly activations are indicated by coloured dashed circles. In step 1, the covariance matrix was computed. Next, eigenvalue decomposition revealed three eigenvalues above the critical value given by the Marčenko-Pastur law (red), indicative of three assembly patterns (step 2). Independent component analysis was used to extract the weight patterns of the 3 assemblies (step 3). Neurons with high weight in each of the patterns match the original assembly neurons. Finally, projecting the weight patterns back onto the spike matrix allows the reconstruction of assembly activation time courses (step 5).

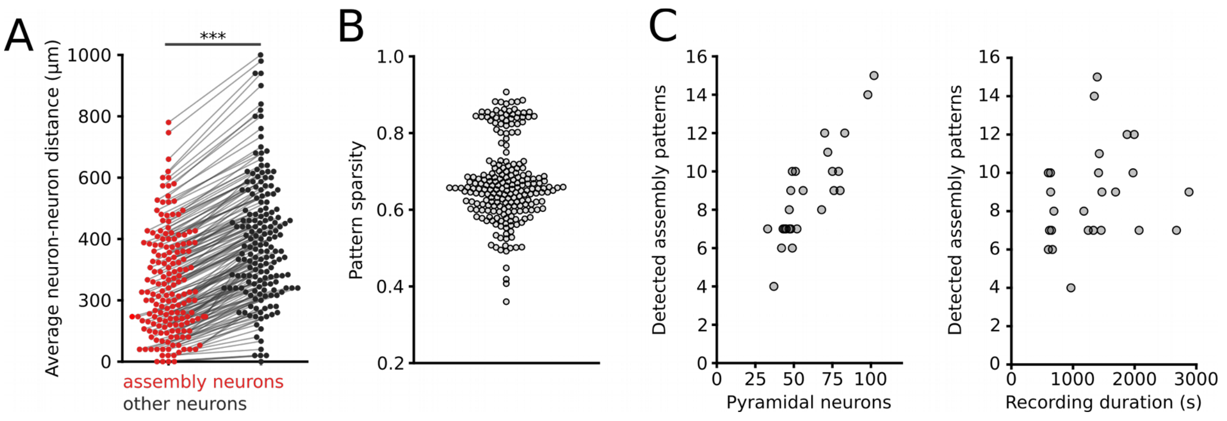


**Supplementary Fig. 3: Assembly properties.**

A: Assemblies are spatially clustered. Intersomatic distance for pairs of assembly neurons is significantly smaller than the distance between randomly drawn neurons. n=173 pairs of assembly vs. non-assembly groups from 13 mice. $p=10^{-11}$, paired *t*-test. B: Average pattern sparsity. High sparsity values indicate that the patterns are dominated by high weights of few neurons. C: The number of detected assembly patterns depended on the number of simultaneously recorded pyramidal neurons (left, Spearman's r = 0.820, $p=5*10^{-7}$) but not on recording duration (right, r = 0.293, p=0.156). 25 sessions from 13 mice.

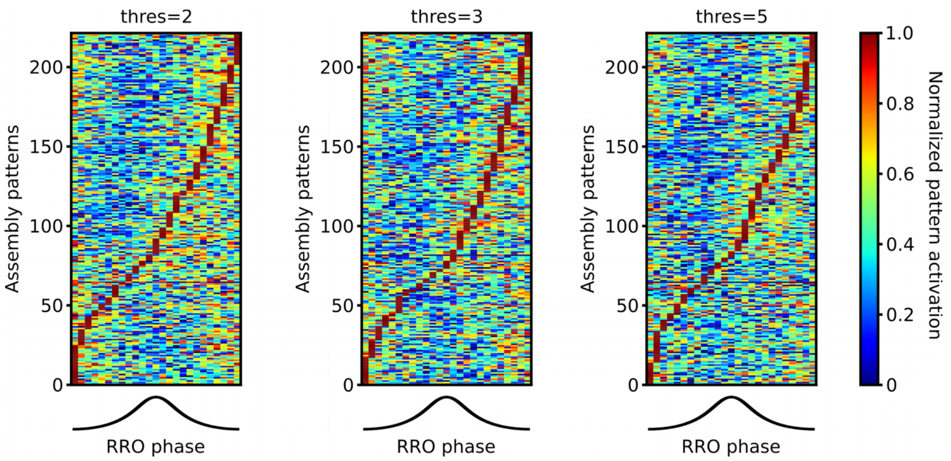


**Supplementary Fig. 4: Assemblies activate during the descending phase of RRO.** Concentration of assembly activations during the descending phase is observed irrespective of the detection threshold. Data show peak normalized assembly activations for threshold values of 2, 3 and 5 as a function of RRO phase. Data are from 25 sessions from 13 mice.

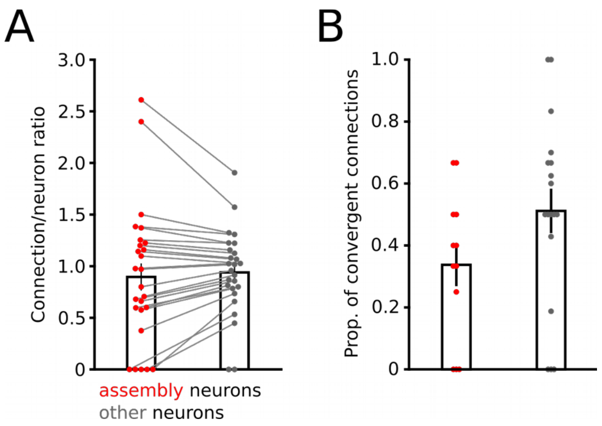


**Supplementary Fig. 5: Properties of assembly- and non-assembly-interneuron connections.** A: Comparable connections/neurons ratio for outgoing connections from assembly and non-assembly neurons onto local putative interneurons indicates similar connection probability. n=25 sessions from 13 mice, p=0.806, paired *t*-test. B: Similar proportion of convergent connections (i.e. with more than one presynaptic pyramidal cell impinging on a postsynaptic interneuron). 12 sessions from 10 mice, p=0.108, *t*-test. Bars show mean ± sem.
